# Dietary phenolics and their microbial metabolites are poor inhibitors of trimethylamine oxidation to trimethylamine N-oxide by hepatic flavin monooxygenase 3

**DOI:** 10.1101/2023.04.21.537826

**Authors:** Lisard Iglesias-Carres, Sydney A. Chadwick-Corbin, Michael G. Sweet, Andrew P. Neilson

## Abstract

High circulating levels of trimethylamine N-oxide (TMAO) have been associated with cardiovascular disease risk. TMAO is formed through a microbiome-host pathway utilizing primarily dietary choline as a substrate. Specific gut microbiota transform choline into trimethylamine (TMA), and, when absorbed, host hepatic flavin-containing monooxygenase 3 (FMO3) oxidizes TMA into TMAO. Chlorogenic acid and its metabolites reduce microbial TMA production *in vitro*. However, little is known regarding the potential for chlorogenic acid and its bioavailable metabolites to inhibit the last step: hepatic conversion of TMA to TMAO. We developed a screening methodology to study FMO3-catalyzed production of TMAO from TMA. HepG2 cells were unable to oxidize TMA into TMAO due to their lack of FMO3 expression. Although Hepa-1 cells did express FMO3 when pre-treated with TMA and NADPH, they lacked enzymatic activity to produce TMAO. Rat hepatic microsomes contained active FMO3. Optimal reaction conditions were: 50 µM TMA, 0.2 mM NADPH and 33 µL microsomes/mL reaction. Methimazole (a known FMO3 competitive substrate) at 200 µM effectively reduced FMO3-catalyzed conversion of TMA to TMAO. However, bioavailable chlorogenic acid metabolites did not generally inhibit FMO3 at physiological (1 µM) nor supra-physiological (50 µM) doses. Thus, the effects of chlorogenic acid in regulating TMAO levels *in vivo* are unlikely to occur through direct FMO3 enzyme inhibition. Potential effects on FMO3 expression remain unknown. Intestinal inhibition of TMA production and/or absorption are thus likely their primary mechanisms of action.

## 1. Introduction

Cardiovascular disease (CVD) is one of the major causes of death worldwide [1]. A key factor in the development of CVD, cardiovascular risk and mortality is atherosclerosis [2]. In recent years, trimethylamine *N*-oxide (TMAO) has been suggested to be a novel, relevant biomarker and risk factor for the development of atherosclerosis, CVD and cardiovascular mortality [2–4]. However, it is important to highlight that the involvement of TMAO in CVD is still controversial, as some studies do not show a causal relationship between TAMO levels and CVD [5,6]. Overall, controlling circulating levels of TMAO could potentially have a significant benefit for public health.

TMAO is produced through a gut microbiota-host metabolic axis. First, choline is converted into trimethylamine (TMA) by specific gut bacteria with the choline utilization gene cluster *cutC/D*, encoding for TMA-lyases [7,8]. Other quaternary amines, such as L-carnitine and betaine can be converted into TMA by different enzymes as well [2,7,9]. Once TMA is formed in the gut, it can be absorbed by the host and reach the liver where flavin-containing monooxygenases (FMOs), particularly FMO3, oxidize it into TMAO [10–12]. Several studies point out that, once in circulation, TMAO exerts several biological effects that ultimately contribute to atherosclerosis development and a higher risk of cardiovascular mortality [2,9,13]. However, some other studies report that TMAO is not associated with cardiometabolic phenotypes and inflammatory markers in children and adults [5].

Plant phytochemicals, and phenolic compounds in particular, have broadly been studied for their cardioprotective effects [14–16]. However, the mechanisms of this protection remain incompletely understood. In recent years, the ability of phenolics to modulate production of TMA and/or TMAO has emerged as another potential cardioprotective mechanism of action [9]. To explore this possibility, we recently screened and identified several phenolic compounds able to inhibit TMA production in an *ex vivo-in vitro* anaerobic colonic microbiota model [17,18]. One of the most potent inhibitors of TMA production in our model is chlorogenic acid. Further research performed by us and other authors also suggests the potential of chlorogenic acid for the modulation of circulating TMAO levels [17,19,20].

Native phenolics from the diet are generally poorly bioavailable, and a large percentage of the ingested dose reaches the colon where it is subjected to gut microbiota fermentation [21,22]. This makes the gut and gut microbiota an appealing target to study the role of phenolics as inhibitors of TMAO production via inhibiting choline metabolism to TMA by the microbiota [9]. However, a percentage of native phenolics and their gut-derived phenolic metabolites can also be absorbed by the host, and reach both the liver and systemic circulation [21,22]. Thus, inhibiting conversion of TMA to TMAO by hepatic FMO3 may also be a complementary mechanism for reducing TMAO producton *in vivo*. In some cases, the bioavailability of gut microbiota-metabolites is much higher than the native compounds [23]. Of note, gut microbiota-derived phenolic metabolites have shown relevant bioactivities in various peripheral tissue models [23–25]. Furthermore, different phytochemicals have been shown to regulate circulating TMAO levels by reducing the gene expression and/or activity of FMO3 [26–31]. Recently, naringin, a flavonoid glucoside from citrus fruits, has been recently found to inhibit FMO3 activity in an *in vitro* model [32]. However, a strong inhibition of FMO3 is not desirable, as this could lead to trimethylaminuria symptoms, which carry a social burden for thos individuals carrying mutation in FMO3 gene [11,33]. Thus, the aim of this study was to evaluate if bioavailable phenolic metabolites are able to inhibit the production of TMAO in the liver via direct inhibition of FMO3. To achieve this, three different *in vitro* screening models were developed and evaluated: human hepatic cells (HepG2), mouse hepatic cells (Hepa-1), and rat hepatic microsomes.

## 2. Materials and methods

### 2.1. Chemicals and reagents

Ammonium formate, ammonia, ethyl bromoacetate, TMA, TMA-^13^C_3_-^15^N, TMAO, TMAO-d_9_, 3-(4,5-dimethylthiazol-2-yl)-2,5-diphenyl tetrazolium bromide (MTT reagent), DMSO, NADPH, hepatic microsomes from female Sprague-Dawley rats (M9191), quinic acid, chlorogenic acid, caffeic acid, ferulic acid, isoferulic acid, dihydrocaffeic acid, dihydroferulic acid, dihydroisoferulic acid, *p*-coumaric acid, *m*-coumaric acid and methzimazole (MTM) were purchased from Sigma-Aldrich/Millipore (St. Louis, MO, USA). Acetonitrile and water (HPLC grade) were purchased from VWR International (Suwanee, GA, USA). HepG2 and Hepa-1c1c7 (Hepa-1, from a female mouse) cells were purchased from ATCC (Manassas, VA, USA). Dulbecco’s Modified Essential Media (DMEM), nucleoside-free α-minimal essential medium (MEM-α), non-essential amino acids (NEAA), penicillin, streptomycin and gentamicin were purchased from Lonza (Morristown, NJ, USA). Heat-inactivated fetal bovine serum (FBS) was purchased from R&D systems (Minneapolis, MN, USA). HEPES was purchased from Fisher Scientific (Waltham, MA, USA)

### 2.2. General cell culture

HepG2 (human hepatocyte) cells were cultured in growth media consisting of DMEM, 10 % FBS, 0.5 % penicillin/streptomycin, 0.1 % gentamicin, and 1 % NEAA. Cells were maintained at incubator conditions of 37 °C in a 5 % CO_2_/air with 90 % humidity. Growth medium was changed every 2 days, and cells were sub-cultivated 1 – 2 times a week. Subcultivation and treatments were only performed to cells that reached > 80 % confluence. Hepa-1 cell culture followed the same procedure, but grown in MEM-α with 10 % FBS, 1% HEPES, 0.5 % penicillin/streptomycin, 0.1 % gentamicin, and 1 % NEAA.

### 2.3. Screening method in human hepatic cells

#### 2.3.1. TMA toxicity in HepG2

A first set of experiments was performed in HepG2 cells to evaluate potential TMA toxicity. 48 h prior to treatments, cells were seeded at 3.5x10^5^ cells/mL in 24-well plates with 1 mL of growth media. The day of the experiment, growth media was removed and cells were washed once with pre-warmed PBS 1X. Cells were then treated with 950 µL DMEM and 50 µL TMA for final TMA concentrations of 10 – 200 µM. A control without TMA was achieved by treating cells with 950 µL DMEM and 50 µL PBS. Cells were exposed to experimental conditions for 10 h, and toxicity (loss of viability) was assessed via the MTT assay.

#### 2.3.2. TMAO production in HepG2

A second set of experiments was performed in HepG2 cells to evaluate their capacity to produce TMAO. 48 h prior to the treatments, cells were seeded as described above in 24-well plates. 24 h prior to treatment, cells were pre-treated with 100 µM TMA in growth media. The day of the experiment, media was removed and cells were washed once with pre-warmed PBS. Cells were then treated with ± 100 µM TMA and ± NADPH (0 – 40 mM) in DMEM. Toxicity controls included TMA and NADPH-free conditions, and NADPH-free conditions. To assess spontaneous chemical (non-cellular) production of TMAO, cell-free controls with 100 µM TMA and 0 or 2 mM NADPH were included. At selected time points (0, 1.5, 3, 4.5, 6, 7.5, 9, 10.5 and 24 h), aliquots (75 µL) of treatment media were taken, immediately mixed with equal volume of acetonitrile and frozen until TMA and TMAO analyses. At the end of the kinetic study (24 h), all treatments (except cell-free controls) were subjected to cytotoxic evaluation by MTT assay.

### 2.4. Screening method in Hepa-1 cells

Due to the lack of observed TMAO production in HepG2, we next investigated the potential of Hepa-1 (rat hepatocyte) cells as a alternative *in vitro* cell model for TMAO production due to their previously reported FMO activity [34]. 48 h prior to the treatments, cells were seeded as described above in 24-well plates. 24 h prior to treatment, cells were pre-treated with 100 µM TMA in growth media. The day of the experiment, media was removed and cells were washed once with pre-warmed PBS. Cells were then treated TMA (0 – 100 µM) and NADPH (0.0 – 2 mM) in MEM-α or complete growth media. A batch of experiments with the same TMA and NADPH treatments concentrations were performed in Hepa-1 cells, but without TMA pretreatment. None of these non-pretreatment conditions reported TMAO production (data not shown). At time 0 h and 24 h aliquots (100 µL) of treatment media were taken, immediately mixed with an equal volume of acetonitrile and frozen until TMA and TMAO analyses.

### 2.5. Toxicity assessment in cell lines

Potential cytotoxic effects of treatments on cells were evaluated through the MTT assay [17]. At the end of the kinetic studies, remaining treatment media was removed and cells washed with 1 mL PBS. Cells were then treated with 200 µL DMEM (HepG2) or MEM-α (Hepa-1) supplemented with 0.5 mg/mL MTT reagent. After incubation at 37 °C for 20 min, media was aspirated and 1 mL DMSO was added and shaken until reduced MTT crystals were solubilized. An aliquot of 100 µL was read at 560 nm in a SpecraMax iD3 plate reader (Molecular Devices, San Jose, CA, USA). Results were relativized to control conditions, and expressed as a percentage of change against control (untreated cells) ± SEM (*n*=3).

### 2.6. Screening method in microsomes

A rat microsome-based screening method was developed to evaluate the potential of bioavailable phenolics and their microbial metabolites to inhibit FMO3-catalyzed TMAO formation. Microsome assays were conducted in phosphate buffer (pH 7.4) in a water bath at 37 °C. Different concentrations of microsomes (0 – 200 μL/mL reaction; 17.06 ± 0.24 mg protein/mL), TMA (0 – 50 μM) and NADPH (0 – 4 mM), and reaction times (0-45 min) were studied to optimize the assay. Further, MTM (a known FMO3 competitive substrate [35,36]) was evaluated 0 – 1 mM. Quinic acid, chlorogenic acid, caffeic acid, ferulic acid, isoferulic acid, dihydrocaffeic acid, dihydroferulic acid, dihydroisoferulic acid, *p*-coumaric acid and *m*-coumaric acid were used as representative bioavailable metabolites of chlorogenic acid [16] and evaluated as TMAO production inhibitors at a physiologically relevant dose (1 μM), as well as a pharmacological dose (50 μM). At selected timepoints (0 – 45 min), reaction aliquots (25 – 50 μL) were collected and mixed with ice-cold acetonitrile. Then, samples were filtered through a AcroprepAdv 0.2 μm WWPTFE 96-well filtering plates (Pall Corporation, Port Washington, NY, USA) by centrifugation (10 min, 3,400 x g), collected in a fresh 96-well collection plate and frozen at –80 °C until extraction/derivatization with an OT-2 liquid handler robot (Opentrons, NY, USA).

### 2.7. Extraction and quantification of TMA and TMAO

Extraction and quantification of TMA and TMAO from cell culture and microsome reaction supernatants were carried out as described by Iglesias-Carres *et al*. [17]. To extract TMAO, 25 μL of sample:acetonitrile mixture (50:50; v:v) were mixed with 10 μL of ZnSO_4_ (5 % w/v in water), 100 μL acetonitrile and 20 μL TMAO-d_9_ [internal standard (IS); 5 μM] in 96-well plates. After sonication for 5 min in a water bath, samples were filtered through AcroprepAdv 0.2 μm WWPTFE 96-well filtering plates (Pall Corporation, Port Washington, NY, USA) by centrifugation (10 min, 3,400 x g), collected in a fresh 96-well collection plate and frozen at –80 °C until UHPLC-MS/MS analysis. TMA requires derivatization to ethyl betaine to facilitate LC-MS/MS ionization. Briefly, 25 μL of cell supernatant:acetonitrile (50:50; v:v) were mixed with 20 μL of TMA-^13^C_3_-^15^N IS solution (5 μM), 8 μL concentrated ammonia and 120 μL ethyl bromoacetate (20 mg/mL), and let sit for 30 min. Then, 120 μL 50 % acetonitrile/0.025 % formic acid in distilled water were added. Samples were then filtered and stored as described above. Sample and reagent plating were performed using an OT-2 liquid handler (Opentrons, NY, USA).

After extraction, analytes were analyzed via UPLC-ES-MS/MS. TMAO and TMA were analyzed separately, but with the same UHPLC-ESI-MS/MS method employing distinct MS/MS transitions. Briefly, separation was achieved on a Waters Acquity UPLC (Milford, MA, USA) with an ACQUITY BHE HILIC column (1.7 μm, 2.1x100 mm) coupled to an ACQUITY BHE HILIC pre-column (1.7 μm, 2.1x5 mm). Mobile phases were 5 mM ammonium formate in water (pH 3.5) (A) and acetonitrile (B). The gradient was isocratic (80 % B) for 2 min, with a flow rate of 0.65 mL/min. Colum temperature was 30 °C, and autosampler at 10 °C. Quantification was achieved by coupling UPLC with a Waters Acquity triple quadrupole mass spectrometer. Source and capillary temperatures were 150 and 400 °C, respectively. Capillary voltage was 0.60 kV, and desolvation and cone gas flows (both N_2_) were 800 and 20 L/h, respectively. Electrospray ionization (ESI) was operated in positive mode, and data were acquired by multiple reaction monitoring (MRM) in MS/MS mode. Multi-reaction monitoring (MRM) fragmentation conditions of analytes and IS compounds can be found in **Table S1**.

For quantification, HPLC water was spiked with 0 – 200 µM TMA and TMAO standards to obtain external calibration curves. Samples were quantified by interpolating analyte/IS peak abundance ratios in the standard curves. Data acquisition was carried out using Masslynx software (V4.1 version, Waters). Method sensitivity was determined by limit of detection (LOD) and limit of quantification (LOQ), respectively defined as the concentration of analyte corresponding to 3 and 10 times the signal/noise ratio. LOD and LOQ show the lowest concentration required to be injected (5 µL) for analyte detection and quantification. Method detection limit (MDL) and method quantification limits (MQL), respectively defined as the concentration of analyte corresponding to 3 and 10 times the signal/noise ratio, represent the minimum concentration of analytes in cell supernatant samples to be detected and quantified. Working linearity range concentrations also refer to concentrations of standards on the volumes added to the extraction/derivatization procedure. These method quality parameters can be found in **Table S2**.

### 2.8. FMO3 quantification

FMO3 protein content/presence was evaluated in HepG2, Hepa-1 and rat microsomes. HepG2 were cultured as described above in T25 flasks with or without TMA (50 µM) and NADPH (0.2 mM) for 24 h in triplicates. Hepa-1 followed the same treatments, but with complete media instead of DMEM. Cells were washed twice with PBS, scraped off in 3mL of PBS and collected in 15mL centrifuge tubes. Cells were then centrifuged (5 min, 800 rpm) andreconstituted in 1 mL fresh PBS. Cells were washed twice with 5 mL PBS by centrifgugation and resuspension, and reconstituted in 1 mL fresh PBS. Cell suspensions and microsomes were lysed with1:1 (v:v) RIPA with 5 min sonication in a water bath, centrifuged (10 min, 3,400 x g, 4 °C) and supernantants collected and frozen at – 80 °C until analyses. FMO3 protein content was evaluated by species-specific ELISAs (MyBioSource, San Diego, CA, USA) per kit instructions: mouse-specific FMO3 (kit MBS9327471) for Hepa-1 cells, rat-specific FMO3 (kit MBS9901107) for rat microsomes, and human-specific FMO3 (MBS9901102) for HepG2 cells. Pierce BCA protein assay kit (Thermo Scientific, Rockford, IL, USA) was used for total protein quantification of homogenates used for ELISAs. Results were expressed as µg FMO3/mg protein ± SEM (*n*=3).

### 2.9. Statistics

Prism 8.0 (GraphPad, La Jolla, CA, USA) was used for statistical analyses and graphing. In all cases, statistical significance was defined *a prioi* as *p*<0.05. Specific statistical tests employed varied based on experimental design, and are stated in the caption for each graph.

## 3. Results

### 3.1. TMA and NADPH toxicity on HepG2 cells

We first evaluated if TMA concentrations between 10 – 200 µM were toxic to HepG2 cells in order to determine the maximum concentration that could be used in the assay while maintaining viable cells. No significant changes in cell respiration (MTT) were reported between concentrations after 10 h TMA exposure (Fig. 1A), suggesting TMA concentrations up to 200 µM are not toxic to HepG2 cells. Next, HepG2, cells were pre-treated with 100 µM TMA 24 h prior to treatment with 100 µM TMA and a range of NADPH concentrations (**Fig. 1B**). Despite a few statistically significant differences, co-treatment with TMA 100 µM and different concentrations of NADPH in HepG2 did not change cell respiration more than 20% compared to untreated cells. Our results show an inverse “U-shaped” or hormetic effect of NADPH on cell respiration, suggesting that an optimal viability of cells is obtained at NADPH concentrations of 0.2 mM when co-treated with TMA 100 µM.

**Figure 1:**
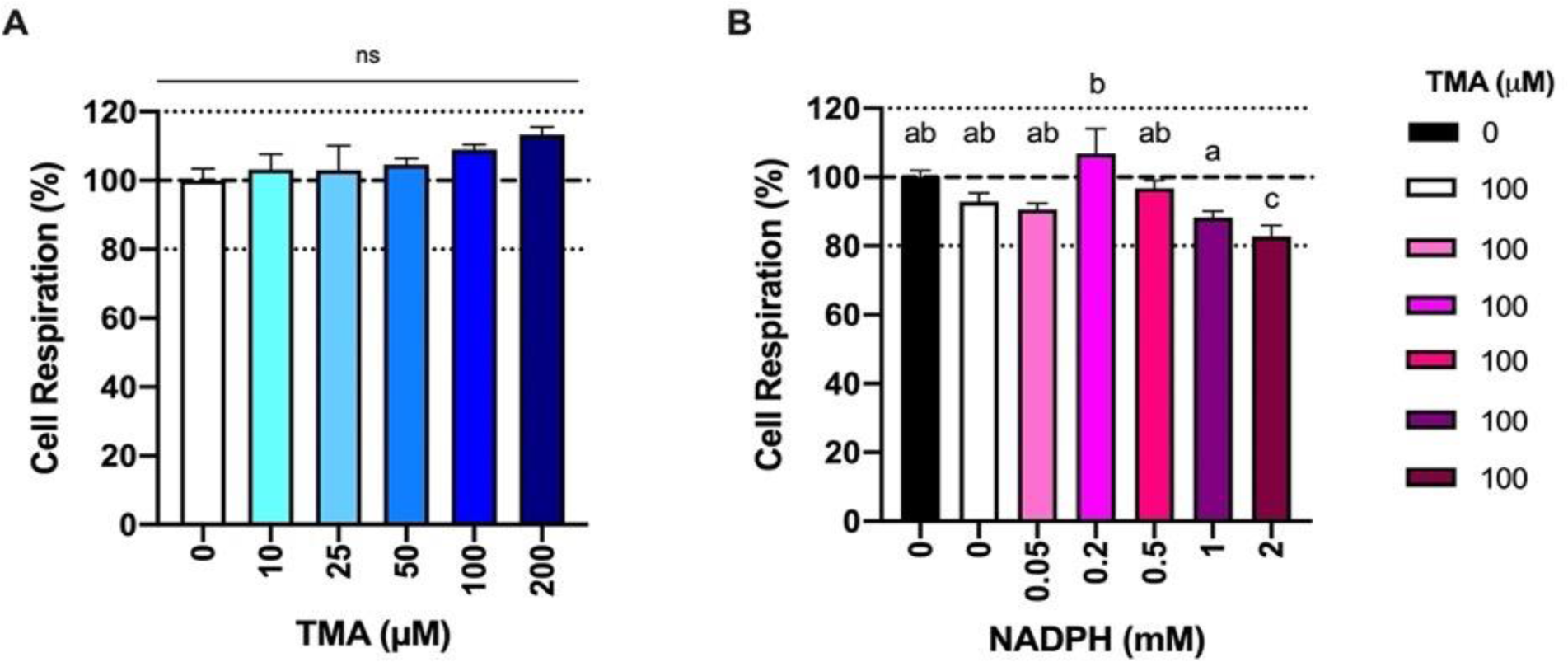
Cell respiration rate (MTT assay) in HepG2 cells exposed to different concentration of TMA (0 – 200 µM) for 10 h (**A**), and HepG2 pre-treated with TMA 100 µM for 24 h and exposed to TMA 100 µM and different concentrations of NADPH (0 – 2 mM) for 24 h (**B**). Results are expressed as percentage of change against control conditions ± SEM (*n*=3). Different letters indicate statistically significant differences (*p*<0.05) by One-Way ANOVA (Tukey’s *post ho*c test).

### 3.2. TMA stability and lack of TMAO production in HepG2 culture

To eliminate the possibility that TMAO was produced by a non-cell catalyzed reaction in our assay, different conditions were evaluated (**Fig. 2A**). In cell-free conditions (red bars) with 100 µM TMA and 2 mM NADPH (highest concentration tested), TMA was stable and its concentrations did not decrease over time. Similar results were reported when neither HepG2 cells nor NADPH were present in the mixture (blue bar). TMAO was not detected in any of the cell-free conditions. When HepG2 were treated for 24 with 100 µM TMA and 2 mM NADPH, TMA levels still did not decrease (black bars), nor was TMAO detected. The full 24 h kinetic plot of TMA in these epxperimental conditions can be found in the form of absolute values in **Figure S1A**, and in the form of percentage against control in **Figure S1B**. This lack of production of TMAO (data not shown) and degradation/use of TMA (**Fig. 2B**) was also reported in HepG2 cells co-treated with TMA 100 µM and NADPH 0 - 2 mM. The full 24 h kinetic plot of TMA in these epxperimental conditions can be found in the form of absolute values in **Figure S2A**, and in the form of percentage against control in **Figure S2B**. Additionally, TMA was also not produced in our first trial (no pre-treatment with TMA nor co-treatment with NADPH) with HepG2 over 10 h (data not shown). Overall, our results show that TMA is not spontaneously chemically degraded nor converted into TMAO in our experimental design in the absence of cells, and that HepG2 are also not capable of transforming TMA into TMAO (likely due to lack of FMO3).

**Figure 2:**
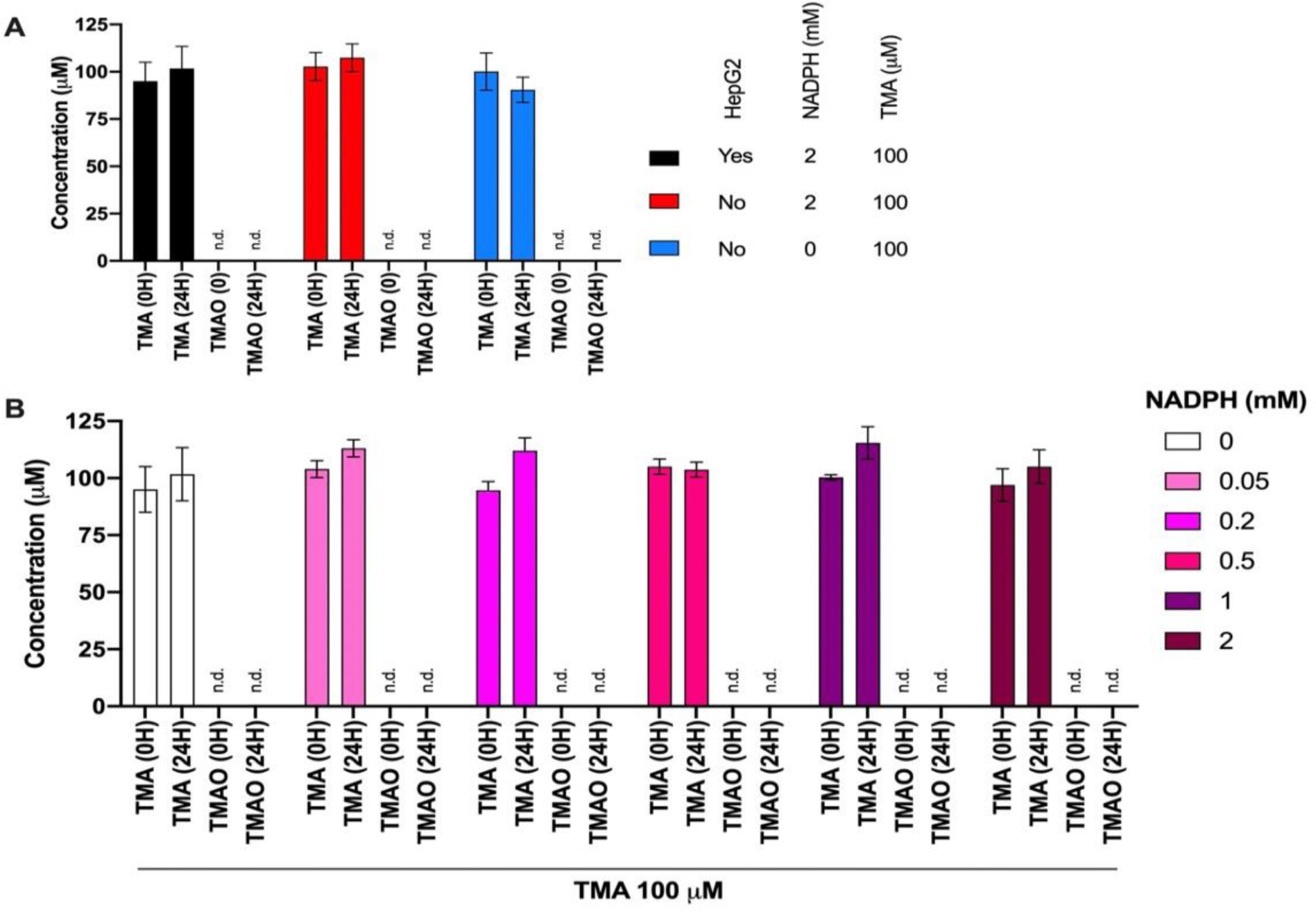
Lack of spontaneous (non cell-catalyzed) production of TMAO in HepG2 cell culture conditions for 24 h (**A**). Lack of TMAO production in HepG2 cells pretreated with 100 µM TAM 24 h prior to treatment with TMA 100 µM and different concentrations of NADPH (0 – 2 mM) for 24 h (**B**). Results are expressed as percentage of change against control conditions ± SEM (*n*=3).

### 3.3. Lack of TMAO production in Hepa-1 cell cultures

Given the lack of TMAO production in human HepG2 cells, we next evaluated mouse Hepa-1 cells as another potential TMAO-producing cell model due to previously reported FMO activity of this cell line towards other FMO substrates [34]. The lack of toxicity and TMA degradation in the HepG2 assays was used to design the approach in Hepa-1 cells. Hepa-1 cells were pre-treated with 100 µM TMA for 24 h before the kinetic experiment to promote FMO3 gene expression and activity. The initial (0h) and final (24h) concnetrations of TMA on pre-treated Hepa-1 cells co-exposed to 10 – 100 µM TMA and 1 mM NADPH were fairily constant (**Fig. 3A**), while TMAO was not detected at 0 or 24 h. Of note, background (no TMA and no NADPH) as well as 0 µM TMA treatments resulted in initial and final TMA concentrations around 7 µM and 11 µM, respectively. This is most likely due to remaining TMA from pre-treatment. Simmilarly, pre-treated Hepa-1 cells co-exposed to 0.2 – 2 mM NADPH and 50 µM TMA reported a slight decrease of TMA over 24h, and a lack of production of TMAO (**Fig. 3B**). In this case, background (no TMA and no NADPH) was the same condition as before, and NADPH 0 mM reported a similar TMA behaviour as + NADPH conditions. Overall, our results indicate that TMA is not being metabolized into TMAO by Hepa-1 cells within 24 h. All these conditions were performed in MEM-α media. We then evaluated if the use of complete growth media would substantially alter TMAO production. **Figure 3C** shows that Hepa-1 cells pre-treated and non-pretreated with TMA are unable to metabolize TMA into TMAO when co-cultured with TMA 50 µM and NADPH 1 mM for 24h. Although cell toxicity (MTT) data is not reported, cells cultured with complete media reported a higher mitochondrial respiration of those cultured with MEM-α, regardless of pre-treatment or non-pretreatmet with TMA. Taken together, our data shows that Hepa-1 cells are also not an appropriate model in which to study TMAO formation.

**Figure 3:**
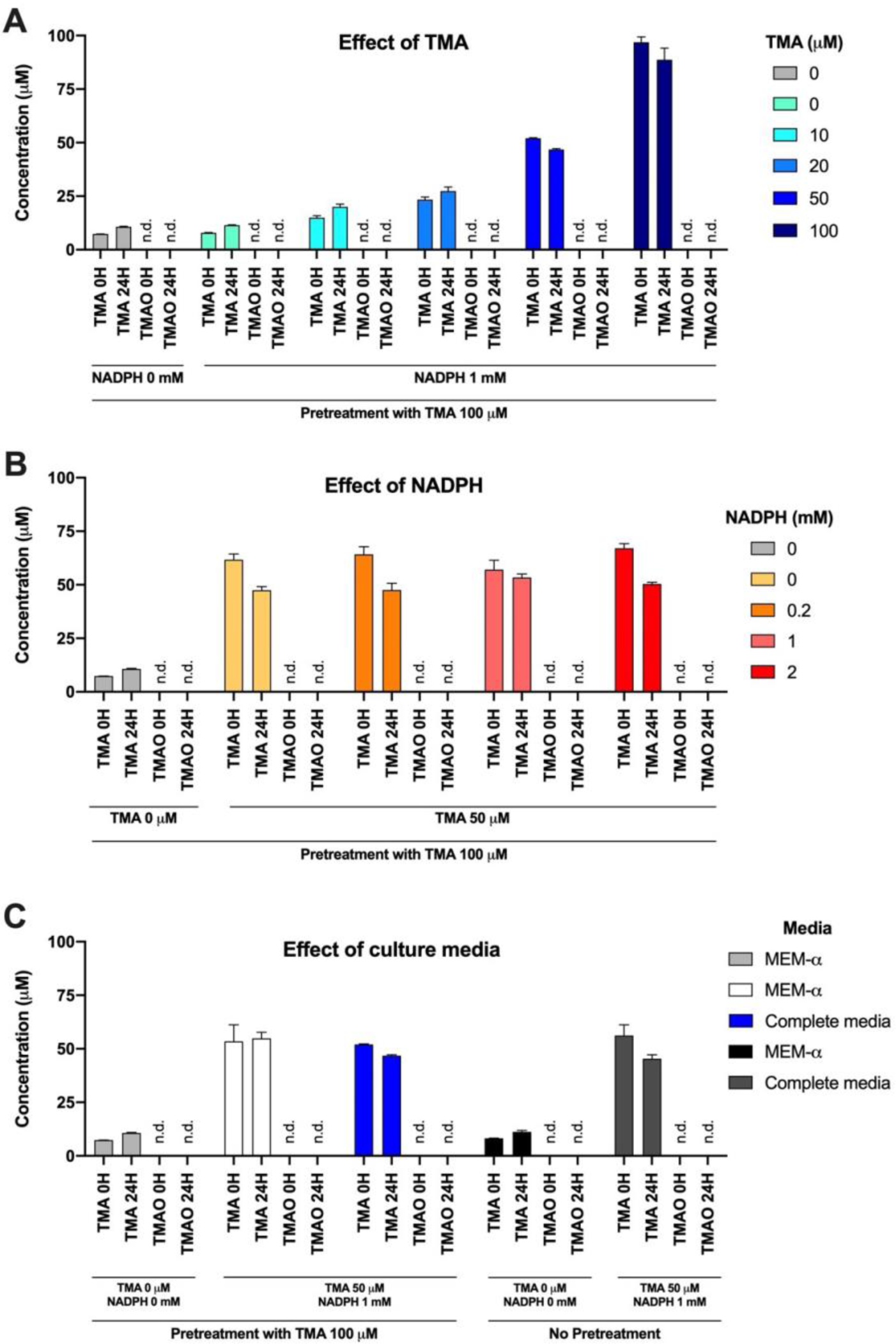
Lack of TMAO production in Hepa-1 cells. Effect of TMA concentration: Hepa-1 cells pre-treated with 100 µM TMA for 24 h, and then co-cultured with TMA 0 – 100 µM and NADPH 1 mM in MEM-α (**A**). Effect of NADPH concentration: Hepa-1 cells pre-treated with 100 µM TMA for 24 h, and then co-cultured with NADPH 0 – 2 mM and TMA 50 µM in MEM-α (**B**). Effect of media: MEM-α and complete (whole) growth media in pre-treated (TMA 100 µM for 24 h) or un-pretreated Hepa-1 cells co-treated with TAM 50 µM and NADPH 1 mM (**C**). Background refers to treatments without TMA and NADPH. Results are expressed as mean ± SEM (*n*=3). TMAO levels were not detected (n.d.) throughout these experimental conditions.

### 3.4. Validation of microsomal production of TMAO

After ruling out HepG2 and Hepa-1 cell lines, we evaluated rat liver microsomes as a potential model. To validate our method, we again evaluated if TMA was stable in our experimental conditions and that it was not oxidized into TMAO by a non-enzymatic catalyzed reaction. To do so, we performed reactions with or without microsomes (100 μL/mL), TMA (10 μM) and/or NADPH (2 mM), as well as 2X these concentrations. Samples were collected at 15 and 30 min (**Fig. 4**). Our results show that the buffer does not contain TMA, and TMAO is not formed in the absence of microsomes. Similarly, microsomes (background, no TMA added) do not supply TMA, and thus TMAO is not formed. This shows that all TMA in our assay is exogenously added, and that all TMAO formed comes from exogenously-added TMA via the microsomnes. We also showed that TMAO is only formed when microsomes and TMA are incubated with NADPH. When no NADPH is added, TMA remains stable and TMAO is not produced. Thus, microsomes, TMA and NADPH are all required factors in the assay. Our data also shows that the reaction is fast: at 15 min all TMA is consumed and converted into TMAO. This is also true when 2 X NADPH (4 mM) and 2 x microsomes (200 μL/mL) are added to the mixture. Finally, our data also demonstrates that NADPH does not spontaneously oxidize TMA into TMAO, and that microsomes are required for the formation of TMAO. Overall, this preliminary assay suggests that 1) rat liver microsomes are a valid model for TMA coinversion to TMAO, and 2) there is no TMA and TMAO contribution of the microsomes and other reagents of the assay; 3) TMAO is only formed when TMA, NADPH and microsomes are co-incubated; and 4) lower concentrations of NADPH and microsomes should be used in further studies in order to capture the kinetics of this reaction.

**Figure 4:**
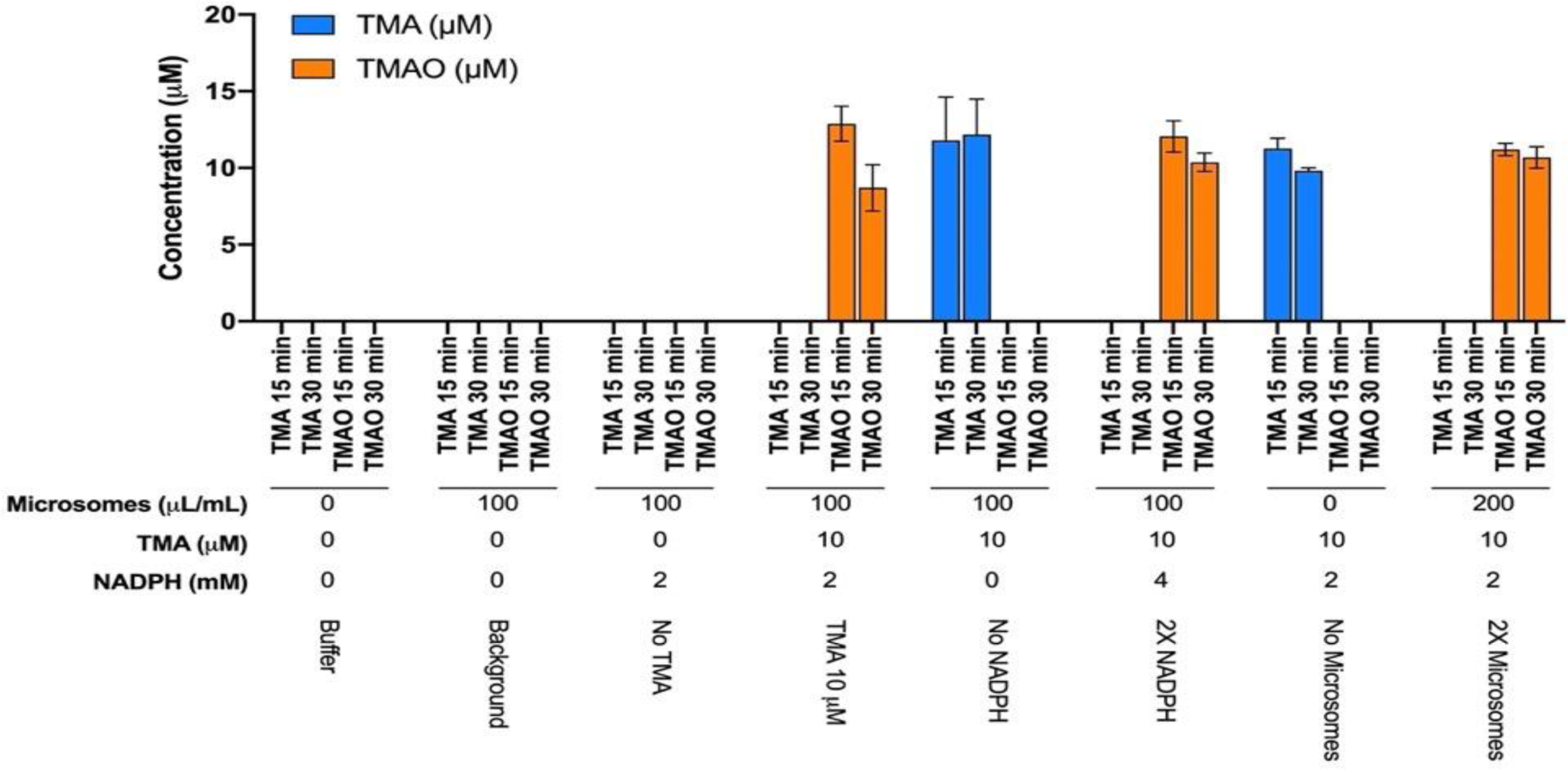
Method validation for TMA and TMAO production and stability in microsome assay. Results are expressed as mean μM ± SEM (*n*=3).

Once our rat liver microsome method was validated as a model of TMA conversion to TMAO, we sought to define appropriate TMA concentrations and characterize the kinetics of the microsomal reaction. Due to the results of our preliminary assay, reactions were conducted at a final concentration of 33 μL/mL microsomes, with 1 mM NADPH. TMA was tested at 0, 20 and 50 μM (**Fig. 5**). Although the quantified concentrations were higher than anticipated based on calculations, our data shows that adding 20 μM TMA (quantified at 29.01 μM at 1 min) was almost completely consumed by 30 min, while when adding 50 μM TMA (quantified at 64.94 μM at 1 min) a significant amount of TMA was still present at 30 min (28.32 μM). Thus, adding 20 μM TMA was selected for further studies. Overall, our selected conditions were 33 μL/mL microsomes, 1 mM NADPH and 20 μM TMA.

**Figure 5:**
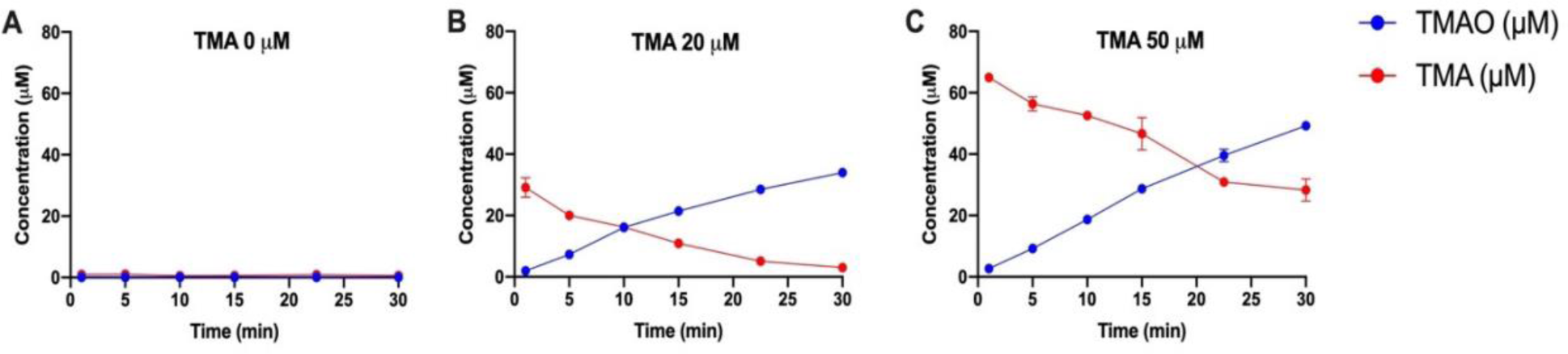
Kinetics of TMA consumption and TMAO production in microsome assay (33 μL/mL) with NADPH 1 mM and TMA 0 μM (**A**), 20 μM (**B**) and 50 μM (**C**). Results are expressed as mean μM ± SEM (*n*=3).

Finally, we evaluated the inhibitory potential of MTM, a known FMO3 competitive substrate [35,36], in our microsome system. This was performed to include a positive control condition in which the conversion of TMA into TMAO would be reduced, as performed by other authors [32]. MTM was tested at 20 – 1000 μM, while other experimental conditions were fixed at 20 μM TMA, 1 mM NADPH and 33 μL/mL microsomes. A background (no TMA and no MTM) condition was included too. Our results corroborated that MTM at concentrations ≥ 50 μM were able to inhibit TMA use and TMAO production (*p*<0.05 by Two-Way ANOVA, main effects: treatment, time; effect of treatment assessed by Tukey’s *post hoc* test) compared to inhibitor-free conditions (**Fig. 6**). However, when Sidak’s *post hoc* test was applied to evaluate the differences at individual time points between inhibitor-free and MTM-supplemented conditions, only concentrations ≥ 200 μM MTM reported relevant statistically-significant changes (**Fig. S3**). Thus, MTM 200 μM was selected as a positive control for further experiments.

**Figure 6:**
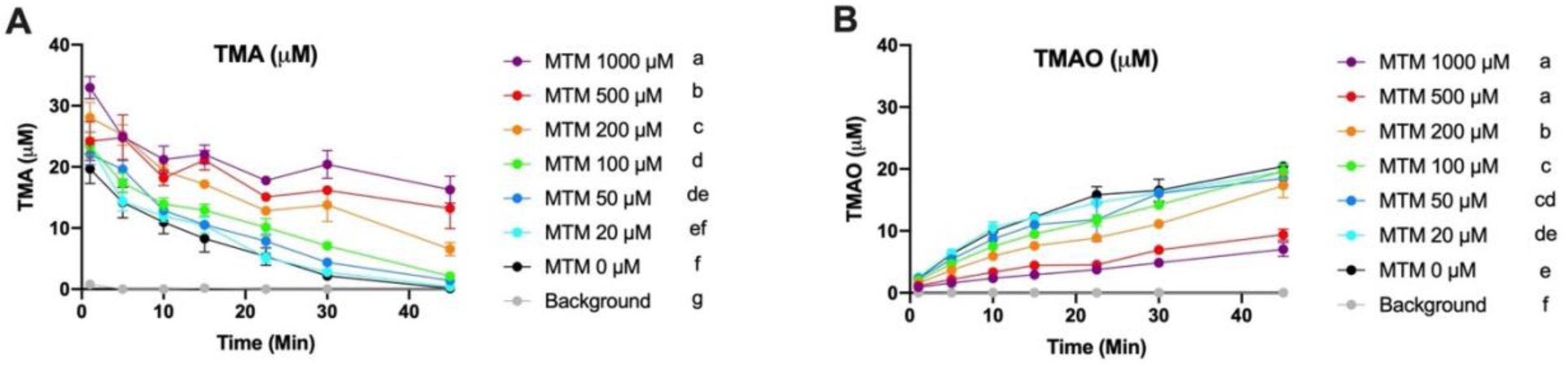
Effect of different concentrations of methimazole (MTM) on TMA consumption (**A**) and TMAO production (**B**) in microsome assay. Results are expressed as mean μM ± SEM (*n*=3). Background corresponds to no TMA or MTM added in the reaction mix. Treatments not sharing common letters (to the right of the legend) have statistically different (*p*<0.05) main effect of treatment by Two-Way ANOVA (Tukey’s *post hoc* test).

### 3.5. Chlorogenic acid and related compounds do not inhibit FMO3 activity

Once our TMAO inhibition assay method was defined and MTM optimized, we next screened for FMO3 inhibitory effects of different chlorogenic acid-related molecules at a physiologically relevant dose of 1 µM. Overall, few of the selected compounds significantly inhibitted TMA use and TMAO production. No treatment other than MTM produced a kinetic curve of TMA use statistically different (overall main effect of treatment by Two-Way ANOVA) from the inhibitor-free control (**Fig. 7A**). The metabolite dihydrocaffeic acid (DHC) was the only compound besides MTM to present a TMAO kinetic curve statistically different from the control (overall main effect of treatment by Two-Way ANOVA, **Fig. 7B**). However, when the AUCs of TMA were analyzed (reflecting total TMA use over time), isoferulic and *p*-coumaric acids (along with MTM) reported statistically significant TMA increases (i.e reduced TMA use) compared to control conditions, suggesting that they inhibited TMA use (**Fig. 7C**). Nevertheless, only DHC (along with MTM) was able to statistically reduce the AUC of TMAO, which agrees with the effects observed in the kinetic curves (**Fig. 7D**). Of note, quinic acid, isoferulic and DHC statistically reduced TMAO percentage against control conditions, while no compound did the same for TMA percentage (**Fig. S4**).

**Figure 7:**
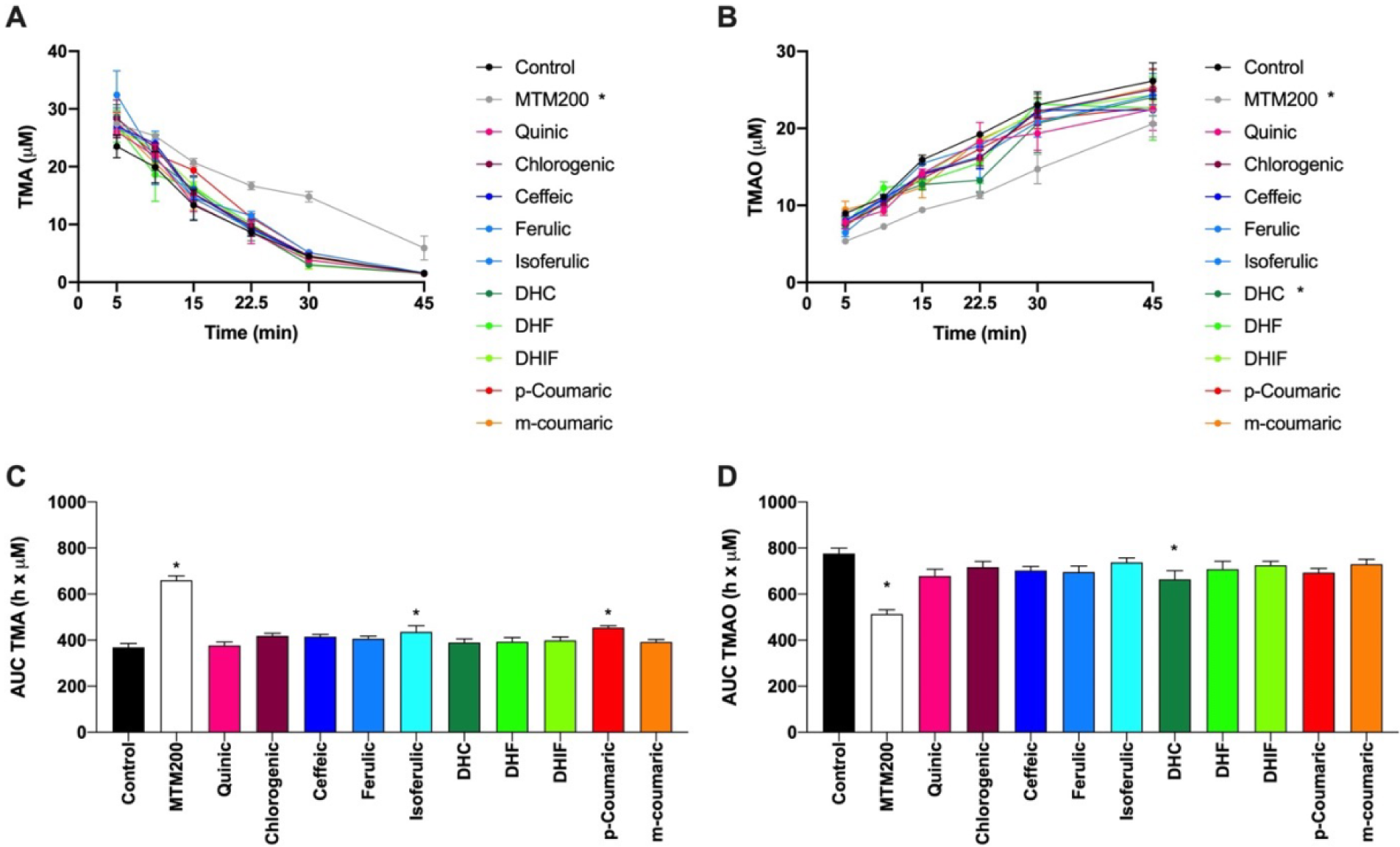
Effect of chlorogenic acid-related compounds (1 µM) in the kinetic curves of TMA (**A**) and TMAO (**B**), as well as the areas under the curve of TMA (**C**) and TMAO (**D**). Results are expressed as mean ± SEM (*n*=3). * indicates statistical differences (*p*<0.05) in the levels of TMA or TMAO against control by Two-Way ANOVA (**A** & **B**) and by One-Way ANOVA (**C** & **D**) (all by Dunnetts’s *post hoc* test).

In order to determine whether higher concentrations would produce inhibition, we evaluated a supraphysiological (pharmacological) dose of 50 µM only obtainable by injection. No compound inhibited the use of TMA at 50 µM. Rather, *p*-coumaric acid and *m*-coumaric acid actually increase production of TMAO, expressed as percentage of change against control conditions (**Fig. S5**). Overall, our data suggests that chlorogenic acid and its related compounds generally are likely to exert a small or no inhibitory effects on TMA conversion to TMAO by FMO3.

### 3.6. FMO3 levels in HepG2, Hepa-1 and rat microsomes

Finally, to further validate our results, levels of FMO3 in the models used were evaluated using species-specific FMO3 ELISA kits (**Fig. 8**). Of note, HepG2 cells, either treated with or without TMA 50 µM and NADPH 0.2 mM, did not report any levels of FMO3 detectable by human FMO3-specific ELISA. On the other hand, Hepa-1 cells reported high levels of FMO3 detectable by the mouse FMO3-specific ELISA after pre-treatment with TMA 50 µM and NADPH 0.2 mM, but their levels were not detected without this pre-treatment. The lack of FMO3 content in HepG2 aligns with the lack of TMAO formation in that model. Interestingly, much greater levels of FMO3 were detected in mouse Hepa-1 cells compared to rat microsomes, although Hepa-1 were unable to produce TMAO and microsomes showed great efficiency to do so.

**Figure 8:**
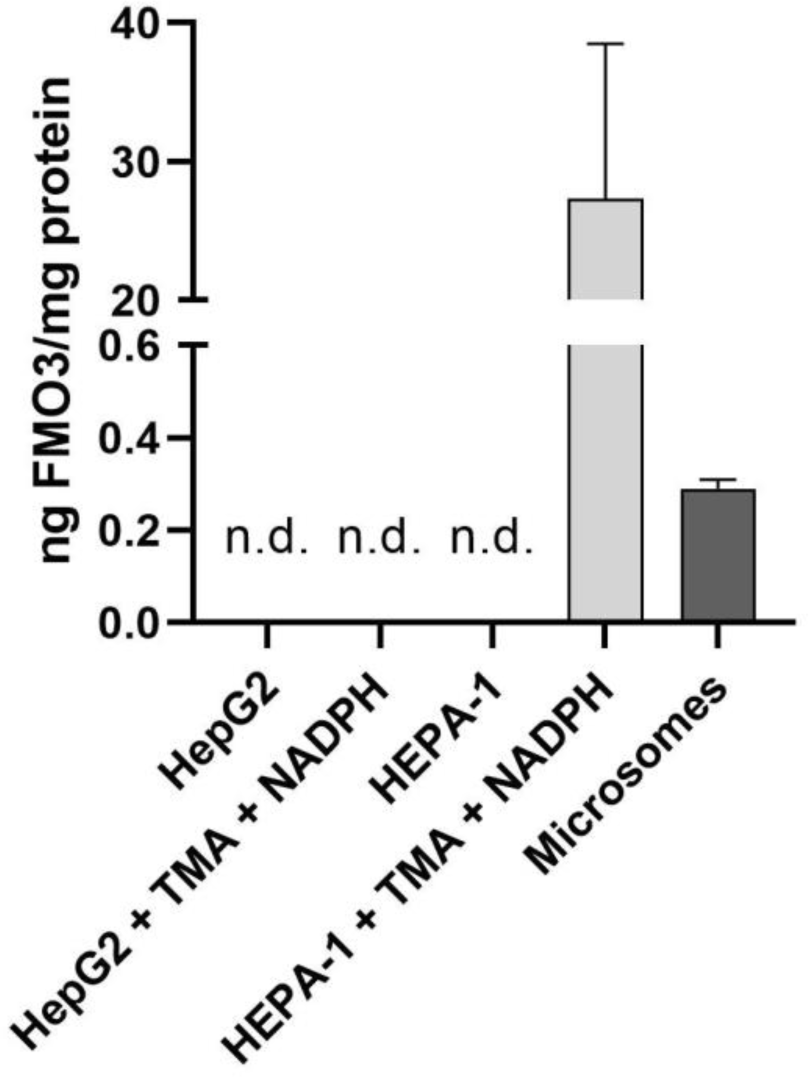
FMO3 levels in HepG2 and Hepa-1 (with and without pre-treatment with TMA 50µM and NADPH 0.2 mM for 24 h) and microsomes (not pre-treated). Results are expressed as ng FMO3 per mg protein ± SEM (*n*=3). Note that the ELISA assay kit employed varied by species: mouse-specific FMO3 (kit MBS9327471) for Hepa-1 cells, rat-specific FMO3 (kit MBS9901107) for rat microsomes, and human-specific FMO3 (MBS9901102) for HepG2 cells.

## 4. Discussion

Cardiovascular disease-related deaths account for 31% of all deaths worldwide [1]. Recently, increased circulating levels of TMAO were shown to promote atherosclerosis development and have been associated with an increase CVD risk [2–4]. However, other studies suggest that high TMAO levels is not associated with a higher CVD risk [5,6]. There is a need for complementary strategies to reduce cardiovascular dicsease burden worldwide. Several phytochemicals, including phenolic compounds, have been shown to inhibit TMAO production through targeting the gut microbiota [9]. We have recently identified several phenolic compounds, such as chlorogenic acid and epicatechin, as potential bioactive compounds to manage TMAO levels by inhibiting microbial production of TMA [17–19]. However, TMA conversion to TMAO is the second step in this metabolic pathway, and it is possible that these compounds could achieve their cardioprotective effects by inhibiting the catalytic activity of FMO3, as reported in previous studies of other phytochemicals [9,32]. We thus aimed to develop an *in vitro* screening method to evaluate the potential of bioavailable phenolic metabolites to inhibit FMO3-catalyzed conversion of TMA to TMAO in the liver. To achieve this, we employed human hepatic cells (HepG2), mouse hepatic cells (Hepa-1), and rat hepatic microsomes.

Firstly, we tried to develop a TMAO inhibition method through the use of HepG2, a convenient and widely used hepatocyte human cell model. HepG2 is an immortalized, commercially-available hepatic cell line isolated from a hepatocellular carcinoma from a 15-year-old white male [37]. Our results showed that concentrations of TMA as high as 200 µM were not toxic to HepG2 cells (**Fig. 1A**). However, HepG2 cells were unable to transform TMA into TMAO. In order to attempt to stimulate the production of TMAO, we pre-treated HepG2 cells with 100 µM for 24 h. FMO3 is a FAD- and a NADPH-dependent enzyme [38]. Thus, after pre-treatment, HepG2 cells were treated with TMA 200 µM and a range of NADPH concentrations from 0 – 2 mM for 24 h. Pre-treatment with TMA and co-treatment with TMA and NADPH was generally not toxic for HepG2 cells (**Fig. 1B**). FMOs are generally thought to be a non-inducible family of genes, although several compounds can increase their gene expression but not their protein content and activity [34]. Pre-treatment with TMA and co-treatment with TMA and NADPH was insufficient to promote HepG2 catalytic conversion of TMA into TMAO (**Fig. 2B**). Previous studies in HepG2 cells have reported FMO activity against different substrates, but not TMA [39]. However, other authors have shown that although the Fmo3 gene is present in HepG2 cells, HepG2 cells are unable to express FMO3 protein [40–42]. When the levels of FMO3 were analysed in lysed cells, FMO3 was not detected (**Fig. 8**). Although the ELISA kit used was not specific to cell clutures, our results agree with the lack of expression of FMO3 reported by others in the literature. To our knowledge, this is the first attempt to develop a FMO3 inhibition method using non-transfected hepatic cells. We have shown that human-derived HepG2 cells are not a good model for our purposes, attributed to their lack of FMO3 expression.

Our second attempt to model FMO3 inhibition (enzme activity, gene expression and/or protein content) was to reproduce our HepG2 approach in a mouse-derived hepatic cell line: Hepa-1. Hepa-1 cells, derived from *Mus musculus* hepatoma, have been previously shown to express the FMO3 gene and possess activity against ethionamide [34]. Thus, Hepa-1 cells were selected to further develop our screening method. Hepa-1 cells were pre-treated with 100 µM TMA for 24 h, and then treated with different concentraions of TMA (0 – 100 µM) and NADPH (0 – 2 mM). Similar to HepG2, Hepa-1 cells were unable to metabolize TMA into TMAO (**Fig. 3A-B**) in MEM-α. Although cell respiration increased when cultured with complete growth media (data not shown), these cells wee unable to produce TMAO when co-treated with TMA 50 µM and NADPH 1 mM (**Fig. 3C**), regardless of pre-treatment with TMA or not. Of note, previous studies have shown that Hep1 cells are able to express FMO3 mRNA but only produce modest amounts of FMO3 [34]. Interestingly, we reported that Hepa-1 cells do express significant levels of some form of FMO3 (detectable by mouse-specific FMO3 ELISA) when pre-treated with TMA 50 µM and NADPH 0.2 mM (**Fig. 8**), albeit with significant inter-sample variability (3 indvidual cell culture flasks were grown up for this assay and analyzed separately). However, this was apparently insufficient for production of TMAO, which indicates that Hepa-1 cells are also not a good model to study FMO3 inhibiton. This was somewhat unexpected, especially when compared to detected FMO3 levels in rat microsomes, an efficient model of TMAO production as per our data. A relevant factor to consider is that our results are relative to the protein content of our samples. Microsomes were much richer in total protein than our Hepa-1 cell lysates (data not shown), and normalization to total protein thus lowered the relative levels of FMO3 compared to Hepa-1 cell lysates. Furthermore, the ELISA kits used to perform these assays were not specific for cells (or microsomes for that matter) and the variability in detected mouse Fmo3 was high. Thus, FMO3 levels should not be directly compared between these models. Rather, this data should be interpreted as a indicator of presence (or lack of) FMO3 in our models. Finally, it is important to note that ELISA detection of a specific antigen and full catalytic activity are not synonymous, and the Hepa-1 cells are not a useful model for studying TMA conversion to TMAO despite high levels of ELISA-detectable FMO3.Overall, our results suggest that although cultured Hepa-1 cells express some form f FMO3, this enzyme is not capable of significant conversion of TMA to TMAO. Overall, our results in immortalized and commercially-available human (HepG2) and mouse (Hepa-1) cell lines highlight the lack of ability of these cell lines to produce TMAO. To our knowledge, although other substrates of FMOs may be metabolized by these cell lines, this is the first study to attempt to assess whether or not these cell lines are able to metabolize TMA into TMAO. The expression of recombinant FMO3 in these or other models or the use of primary hepatic cell cultures could be used to this aim, but not Hepa-1 and HepG2 cells. Noteworthily, this suggests that the study of compounds that can inhibit FMO3 TMAO production should be conducted in other *in vitro* models (*i.e.,* hepatic microsomes) or *in vivo*. While microsomes may provide insights into direct FMO3 enzme inhibitions of chlorogenic acid derived compunds, other mechanisms of actions, such as effects on gene expression and/or protein content regulation, may only be evaluated *in vivo* or in primary hepatocytes [9].

Finally, we aimed to develop a rat microsome screening method that would convert TMA to TMAO and thus allow the identification of direct FMO3 enzyme inhibitors. The presence of FMO3 in rat microsomes is not novel, and has been reported in the past [43]. In our study, female rat microsomes presented a concentration of FMO3 of 0.29 ng FMO3/mg protein (**Fig. 8**), which was much lower compared to that of Hepa-1 cells (27.32 ng FMO3/mg protein). Despite that, these microsomes presented FMO3 activity (**Fig. 4**), which was not the case of Hepa-1. The FMO3 activity in microsomes allowed the development of our methodology. Our initial method validity assays demonstrated that TMA and NADPH need to be present in microsome preparations for TMAO to be formed. The absence of either or both results in a lack of TMAO production (**Fig. 4**). Further, we demonstrated that TMA is stable under our experimental conditions and only oxidized into TMAO with the presence of both microsomes and NADPH. By further optimizing TMA, NADPH and microsomes concentration, we defined our optimized method conditions at a concentration of 20 μM TMA, 33 μL/mL microsomes, 1 mM NADPH and 45 min reaction time (**Fig. 5**). To compare the effects of tretaments (*i.e.,* bioavailable phenolic metabolites) to a positive control (*i.e.,* known FMO3 enzyme inhibitor) we optimized the concentrations of MTM. Results showed that MTM at 200 μM achieved a TMAO production and TMA use inhibition (**Fig. 6** & **Fig. S3**), and thus this concentration of MTM was used in further experiments.

Once our microsomes method was optimized, we evaluated the potential of bioavailable chlorogenic acid metabolites as FMO3 inhibitors (**Fig. 7** & **Fig. S4**) at a physiologically-relevant dose of 1 μM that can be achieved in circulation [16,44]. We selected quinic acid, chlorogenic acid, caffeic acid, ferulic acid, isoferulic acid, dihydrocaffeic acid, dihydroferulic acid, dihydroisoferulic acid, *p*-coumaric acid and *m*-coumaric acid. These compounds are structurally related to chlorogenic acid (**Fig. S6**), and are formed during digestion and/or microbial action on chlorogenic acid. In humans, ∼1/3 of the ingested dose of chlorogenic acid in foods can be absorbed throughout the gastrointestinal tract in its intact, unmetabolized form. About 7% of the ingested dose of chlorogenic acid can be hydrolized in the small intestine into caffeic acid (phenolic moiety) and quinic acid (non-phenolic meoiety), and then absorbed. The remaining chlorogenic acid can reach the colon, where it is fermented by the gut microbiota into dihydrocaffeic acid (reduction of double bond in side chain) as well as smaller phenolic acid derivates. Caffeic and dihydrocaffeic acids can also be absorbed and undergo phase-II metabolism in the gut epithelia and liver through the catalytic activity of catechol O-methyl transferase (COMT). This generates ferulic and isoferulic acids (positional isomers), and its reduced froms dihydroferulic and dihydroisoferulic acids [16]. Thus, these compounds could all be found in circulation following chlorogenic acid consumption. After the consumption of a decaffeinated green coffee beverage rich in chlorogenic acid and related compounds, the maximum concentration in human plasma of the most bioavailable quantified compound were of 5.9 ± 4.2 μM chlorogenic acid, and of 0.4 ± 0.03 μM *p*-coumaric acid for the least bioavailable quantified metabolite [44].

Trimethylaminuria, or “fish-odor syndrome”, is a genetic disorder caused by mutations in FMO3 gene which impairs metabolism of TMA to TMAO [11]. In this condition, increased levels of circulating TMA generate severe psychosocial problems related to the excretion of TMA through sweat, breath and urine due to its fishy smell [33]. Thus, inhibition of FMO3 as a mechanism to reduce TMAO levels should be considered with caution. However, this condition is unlikely to be achieved with phenolic compounds as consumption of phytochemicals has not been previously reported to cause trimethylaminuria, as evidenced by Mediterranean diet intervention [45]. Overall, bioavailable metabolites of chlorogenic acid did not have a dramatic effect on TMA use or TMAO production. Rather, results were very subtle. The only compound able to modulate the kinetic curve of TMAO formation was dihydrocaffeic acid, while none did that for the curve of TMA (**Fig. 7A – B**). This effect was translated into a higher AUC of TMAO compared to control conditions (**Fig. 7D**). Surprisingly, the AUC of TMA for reactions with 1 μM of isoferulic acid and *p*-comaric acid was higher than control conditions, suggesting that these compounds prevented TMA use, but not TMAO production. When results were epressed as percentage of change against control conditions (**Fig. S4**), quinic acid (the non-phenolic part of chlorogenic acid), ferulic acid and dihydrocaffeic acid were able to reduce TMAO % production against control conditions.

Our data suggests that dihydrocaffeic acid could play a small role in the inhibition of *in vivo* TMAO formation of chlorogenic acid through a direct enzyme inhibition [20]. This lack of inhbition of FMO3 by bioavailable phenolic metabolites at physiological doses is in line with the lack of decrease in TMOA levels reported by Mediterranean diet, a diet rich in fruits and vegetables, and thus phenolic compounds [45].

To further validate that bioavailable phenolic metabolites from chlorogenic acid do not or very poorly inhibit FMO3 production of TMAO, the selected metabolites were tested in our assay at a pharmacological dose of 50 μM (**Fig. S5**), impossible to achieve through the diet. Although this is an approach with no relevance to dietary intake, it can be useful to determine whether or not these compounds could potentially inhibit TMAO formation at any dose. A lack of bioactivity at these high doses virtually guarantees the lack of bioactivity at physiologically-relevant doses. Our results show that at these high concentrations, phenolic compounds still do not inhibit the use of TMA (expressed as percentage against control conditions). However, *p*-coumaric and *m*-coumaric acid enhanced the production of TMAO (expressed as percentage of change against control conditions). Our results at these higher contentrations (50 µM) do not correspond to the results found at physiological dose (1 µM). Overall, this indicates that bioavailable metabolites from chlorogenic acid are unlikely to produce an *in vivo* direct FMO3 enzyme inhibition.

There are limitations to the present study. FMO1 may also contribute to conversion of TMA to TMAO [46]. We did not assess FMO1 expresion in microsomes. Hepatic FMO1 decreases in rats with age, however the age of the rats from which hepatic microsomes were prepared is unknown. MTM is also an FMO1 competetive substrate [43]. Therefore, the present study examines enzymes that converts TMA to TMAO, but the finding is the same: chlorogenic acid and its microbial metabolites do not significantly inhibit hepatic conversion of TMA to TMAO. Additionally, the cross-reactivity of the ELISA kits we employed is unknown, and other FMO isoforms may have contributed to measured and reported FMO3 protein expression levels.

Through the *in vitro* approaches in this study, we demonstrated that HepG2 and Hepa-1 cell models are not a good model to study the inhibition of FMO3 activity by phenolic metabolites. This was a result of a lack of production of TMAO in these two hepatic models, attribiuted to the lack of FMO3 protein or its activity. However, we developed a fast screening method to evaluate a direct FMO3 enzyme inhibition by bioavailable phenolic metabolies in rat hepatic microsomes, with a detectable content of FMO3 that presented enzymatic activity. In conclusion, our results show that bioavailable metabolites of chlorogenic acid do not have a strong inhibition potential on the second step of TMAO formation. Rather, some minor inhibitory effects were reported for some metabolites. Of note, even a modest inhibition of hepatic FMO3 could complement the inhibition of TMA production by the microbiome *in vivo*. Overall, the lack of viable cell models to study TMAO production highlights the need of *In vivo* experiments to elucidate if different treatments (chlorogenic acid) are able to contribute to TMAO formation inhibition to mechanisms involving FMO3 (*i.e*., downregulation of gene expression and/or protein content).

## Funding

Partial funding for this project was provided by the North Carolina Agricultural Research Service (NCARS) and the Hatch Program of the national Institute of Food and Agriculture (NIFA), U.S. Department of Agriculture. Funding sponsors had no role in study design; the collection, analysis and interpretation of data; in the writing of the report; and in the decision to submit the article for publication

## Notes

### Competing Interest Statement

The authors have declared no competing interest.

### Summary of Updates

Responses to reviewer critiques from peer review at Journal of Nutritional Biochemistry.

