## Supplemental Material for "Dietary phenolics and their microbial metabolites are poor inhibitors of trimethylamine oxidation to trimethylamine N-oxide by hepatic flavin monooxygenase 3"

**for**

**Table S1: MRM for targeted analytes.**

| Compound | RT (min) | MW (g/mol) | Parent $[M+H]^+$ (m/z) | Daughter $[M+H]^+$ (m/z) | CV (V) | CE (eV) |
|---|---|---|---|---|---|---|
| **TMA** [a] | 0.77 | 145.2 | 146.30 | 118.20 | 34 | 16 |
| **TMA-$^{13}C_3$-$^{15}N$** [a] | 0.77 | 149.2 | 150.27 | 122.19 | 34 | 20 |
| **TMAO** | 1.38 | 75.1 | 76.16 | 58.91 | 40 | 10 |
| **TMAO-$d_9$** | 1.38 | 84.1 | 85.22 | 68.10 | 40 | 12 |

[a] Data for ethyl betaine derivatives.

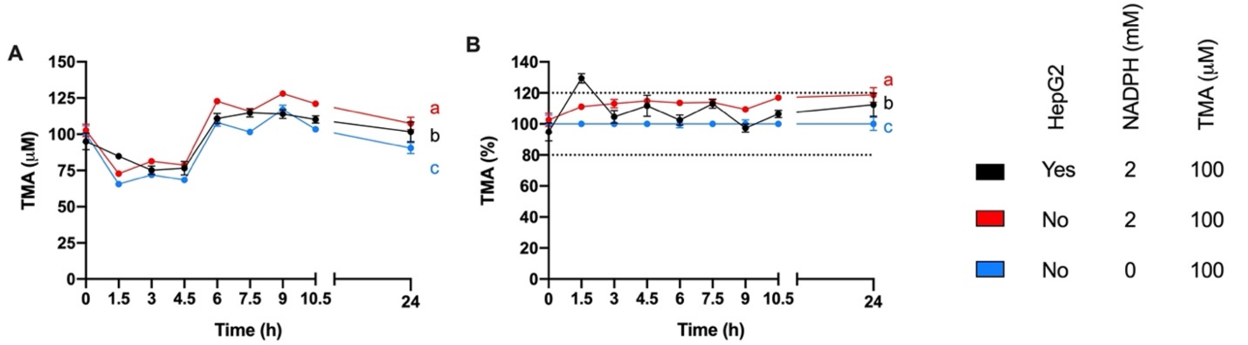


**Figure S1:** Stability of TMA in HepG2 experimental conditions expressed as absolute values (**A**) and percentage against control conditions (**B**). Different letters indicate a statistically significant ($p<0.05$) effect of treatment by Two-Way ANOVA (main effect: treatment; Tukey's *post hoc* test). TMAO levels were not detected (n.d.) throughout these experimental conditions (data not shown).

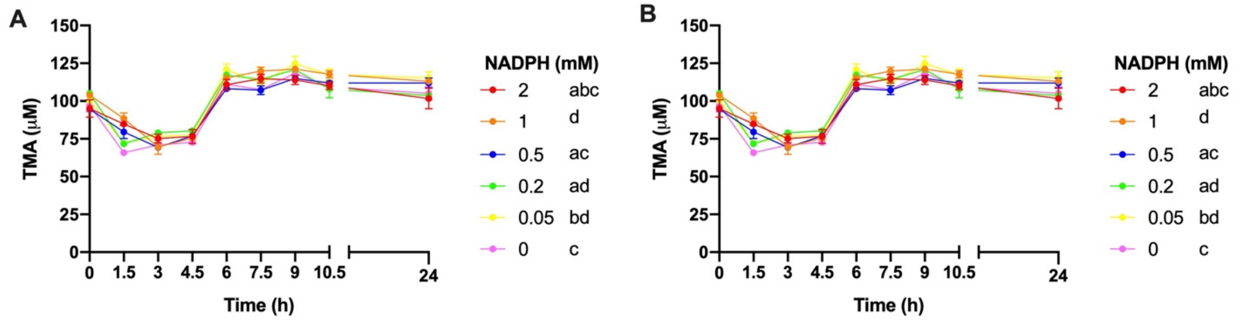


**Figure S2:** Dose-dependence of NADPH on TMA (100 μM) stability expressed as absolute values (**A**) and percentage against control conditions (**B**). Results are expressed as mean ± SEM (*n*=3). Different letters indicate a statistically significant ($p$<0.05) effect of treatment by Two-Way ANOVA (main effects: treatment; Tukey's *post hoc* test). TMAO levels were not detected (n.d.) throughout these experimental conditions (data not shown).

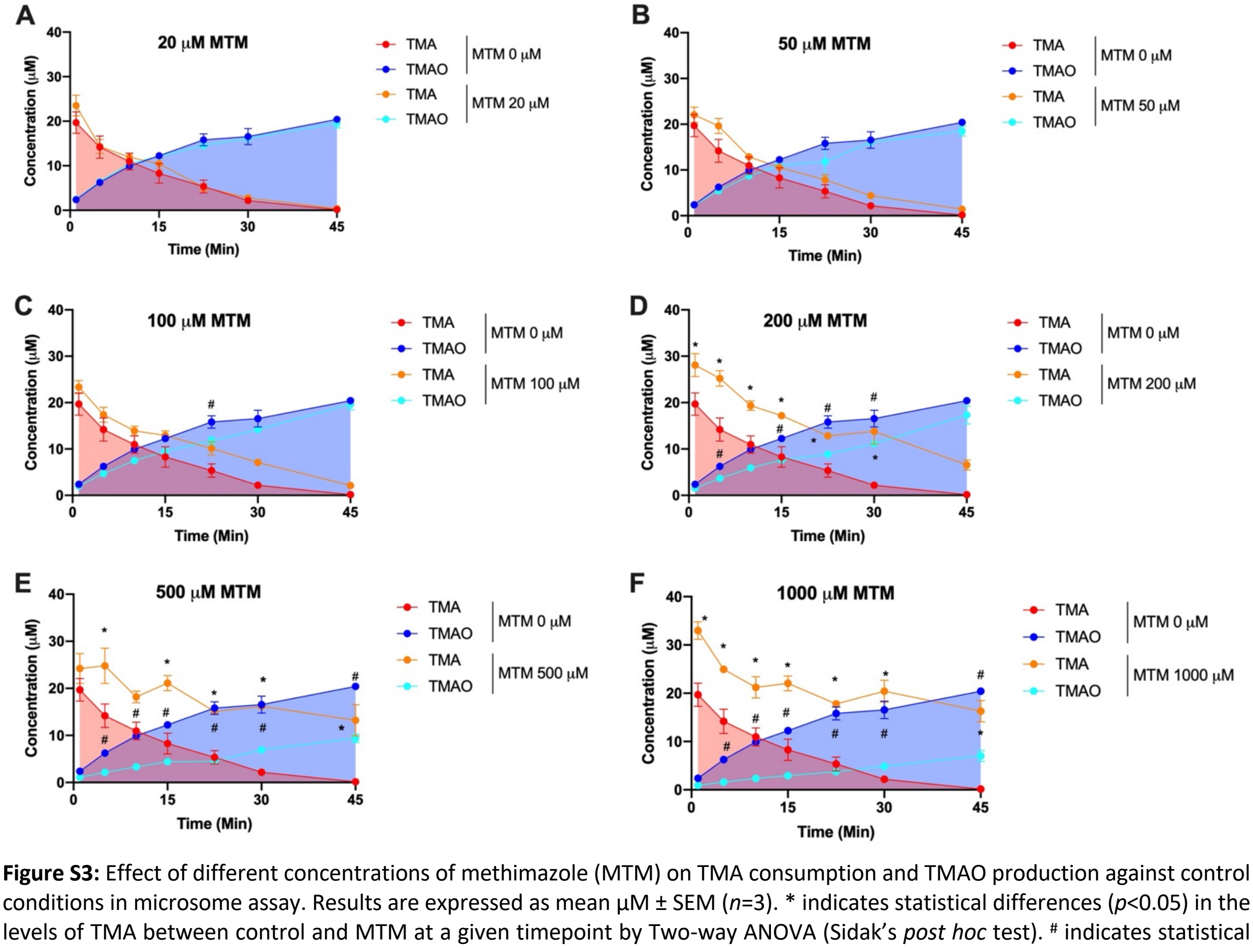


**Figure S3:** Effect of different concentrations of methimazole (MTM) on TMA consumption and TMAO production against control conditions in microsome assay. Results are expressed as mean μM ± SEM (*n*=3). * indicates statistical differences ($p$<0.05) in the levels of TMA between control and MTM at a given timepoint by Two-way ANOVA (Sidak's *post hoc* test). # indicates statistical differences ($p$<0.05) in the levels of TMA between control and MTM at a given timepoint by Two-way ANOVA (Sidak's *post hoc* test).

A

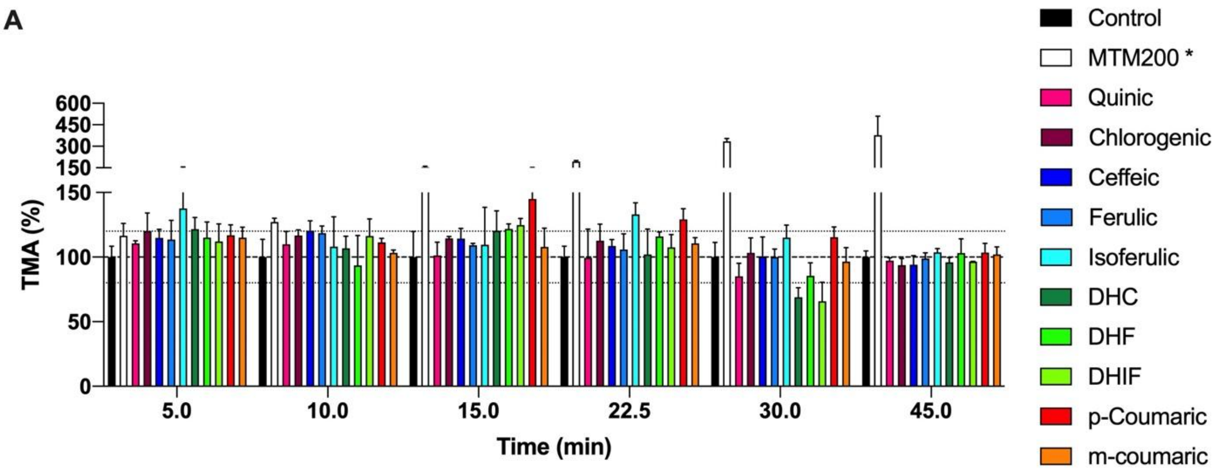


B

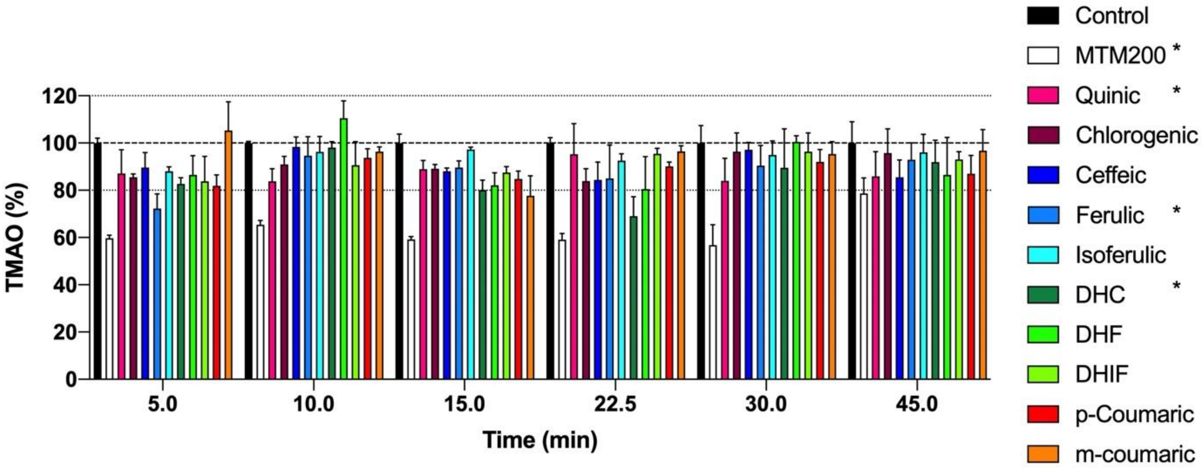


**Figure S4:** Effect of chlorogenic acid-related compounds (1 µM) in the percentage of TMA (**A**) and TMAO (**B**) against control conditions. Results are expressed as mean % ± SEM (*n*=3). * indicates statistical differences ($p$<0.05) in the levels of TMA or TMAO against control by Two-Way ANOVA (**A** & **B**) (Dunnetts's *post hoc* test).

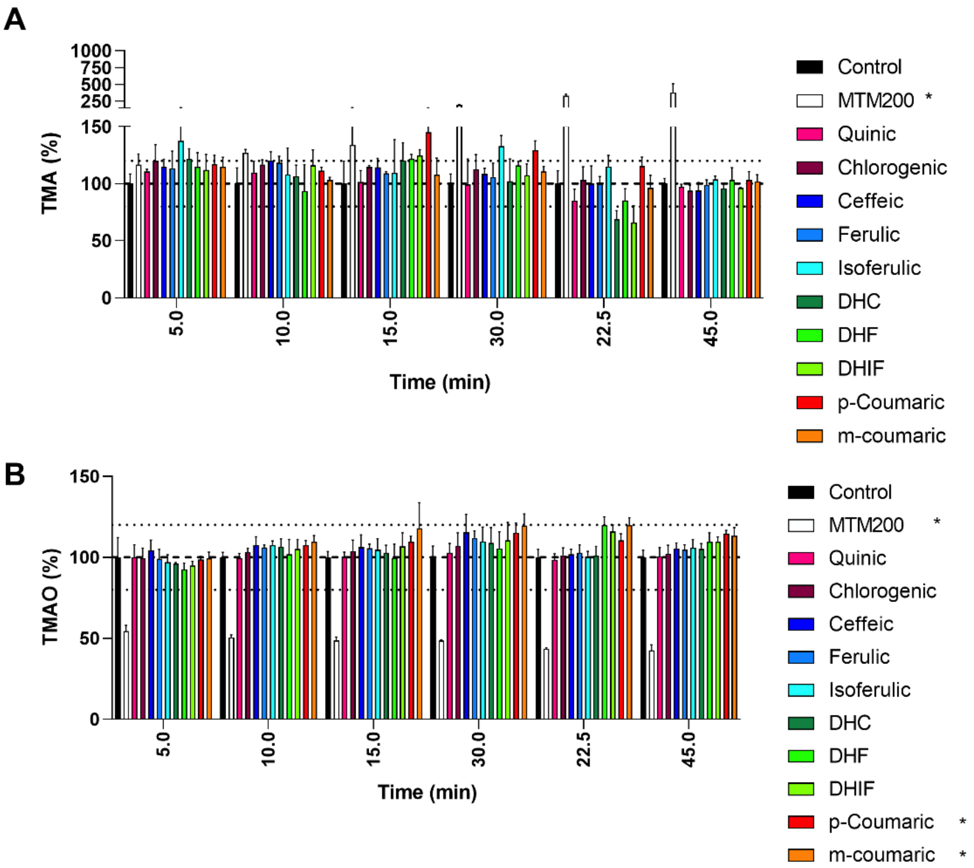


**Figure S5:** Effect of chlorogenic acid-related compounds at supraphysiological doses (50 µM) on the percentage of TMA (**A**) and TMAO (**B**) against control conditions. Results are expressed as mean % ± SEM (*n*=3). * indicates statistical differences ($p<0.05$) in the levels of TMA or TMAO against control by Two-Way ANOVA (**A** & **B**) (Dunnetts's *post hoc* test).

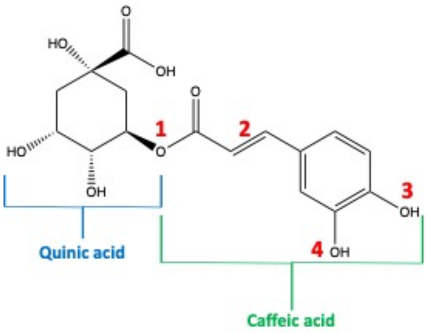


**Structural features of chlorogenic acid and related compounds**

| Compound | Structural features 1 | 2 | 3 | 4 | MW | Other name |
|---|---|---|---|---|---|---|
| **Quinic acid** | Quinic acid | N/A | N/A | N/A | 192.17 | N/A |
| **Chlorogenic acid** | Quinic | = | OH | OH | 354.31 | 5-Caffeoylquinic acid |
| **Caffeic acid** | COOH | = | OH | OH | 180.16 | 3,4-Dihydroxycinnamic acid |
| **Ferulic acid** | COOH | = | OH | OCH3 | 194.18 | 4-Hydroxy-3-methoxycinnamic acid |
| **Isoferulic acid** | COOH | = | OCH3 | OH | 194.18 | 3-Hydroxy-4-methoxycinnamic acid |
| **Dihydrocaffeic acid** | COOH | - | OH | OH | 182.17 | 3'-(3,4-Dihydroxyphenyl)propionic acid |
| **Dihydroferulic acid** | COOH | - | OH | OCH3 | 196.2 | 3'-(4-Hydroxy-3-methoxyphenyl)propionic acid |
| **Dihydroisoferulic acid** | COOH | - | OCH3 | OH | 196.2 | 3'-(3-Hydroxy-4-methoxyphenyl)propionic acid |
| ***p* -Coumaric acid** | COOH | = | OH | H | 164.16 | 4-Hydroxycinnamic aicd |
| ***m* -Coumaric acid** | COOH | = | H | OH | 164.16 | 3-Hydroxycinnamic aicd |

**Figure S6:** Structural features of chlorogenic acid and related compounds used in this study.
